# Efficient Generation and Selection of Virtual Populations in Quantitative Systems Pharmacology Models

**DOI:** 10.1101/028548

**Authors:** R.J. Allen, T.R. Rieger, C.J. Musante

**Affiliations:** Cardiovascular, Metabolic and Endocrine Research Unit, Pfizer Inc., 610 Main Street, Cambridge, Massachusetts, USA.

## Abstract

Quantitative systems pharmacology models mechanistically describe a biological system and the effect of drug treatment on system behavior. Because these models rarely are identifiable from the available data, the uncertainty in physiological parameters may be sampled to create alternative parameterizations of the model, sometimes termed ‘Virtual Patients.’ In order to reproduce the statistics of a clinical population, Virtual Patients are often weighted to form a Virtual Population that reflects the baseline characteristics of the clinical cohort. Here we introduce a novel technique to efficiently generate Virtual Patients and, from this ensemble, demonstrate how to select a Virtual Population that matches the observed data without the need for weighting. This approach improves confidence in model predictions by mitigating the risk that spurious Virtual Patients become over-represented in Virtual Populations.

## Introduction

Quantitative systems pharmacology (QSP) models are an effective approach for gaining mechanistic insight into the complex dynamics of biological systems in response to drug treatment.^1-3^ QSP models in the drug discovery and development process have been utilized for increased confidence in rationale for early development targets, preclinical to clinical translation, and predictions of clinical response to novel therapeutics. To be fit for purpose, these models must include sufficient biological scope and mechanistic detail to link pathway modulation to overall system response^4-8^. Due to the complexity of the biology, the iterative model-building process frequently results in a model that is a large, nonlinear, multi-scale system of equations. Many different data sources are required to quantify QSP models, including *in vitro* and *in vivo* preclinical and clinical data; moreover, the resulting models are frequently under-constrained by any one data set^9^. Therefore, to explore the impact of known variability and uncertainty^10-14^, QSP models are simulated using ensembles of parameterizations often termed “Virtual Patients” or “VPs.” A Virtual Population that reflects individual subject and population level characteristics of a typical clinical cohort provides increased confidence that prospective simulations of response to novel therapeutics will reflect the inter-subject variability seen in the clinic, and may help to identify responders and non-responders to treatment.

Ensembles of VPs are often sufficient for exploring the broad range of responses that are possible from perturbing a model (pharmacologically or otherwise), but the outcome will not necessarily reflect the distribution (e.g., log-normal) of population level data^15^. The result is a range of predictions from the model, which are all possible outcomes but fail to provide insight into the probability of observing that outcome in a clinical trial. Previous authors have overcome this critique by weighting model outputs^13^, or model components^14^, to create Virtual Populations (“VPops”).

Klinke et al. (2008) proposed linearly weighting each VP, with some receiving a weight greater than 1/N (where N is the number of VPs in the ensemble) so that the mean and standard deviation of the VPop match the desired population characteristics^13^. This approach is intuitive and easily implemented, but this also can be computationally expensive, requires re-fitting the VPop each time VPs are added or removed from the analysis, and can result in a dramatic over-weighting of a few select VPs, which may skew the final simulation results. Schmidt et. al. (2013) refined this approach by taking the weights off of the individual VPs and placing them on ‘mechanistic axes.’^14^ Their approach is computationally faster, allows new VPs to be incorporated into the VPop without refitting, and should avoid the problem of overweighting small numbers of VPs. However, their approach is not intuitive to understand or communicate and requires collecting parameters into mechanistic axes that may not explore all the variability and degeneracy in the model appropriately.

Here we propose a new algorithm for generating biologically reasonable VPops. We will show how this algorithm compliments previous approaches by being intuitive, computationally efficient, and avoiding the problem of overweighting VPs. We demonstrate the utility of this new algorithm by fitting the joint distribution of low-density and high-density lipoproteins (LDL and HDL, respectively) from the National Health and Nutrition Examination Survey (NHANES)^16^ to a previously published model of lipoprotein metabolism^17^.

### Method

A flow diagram of the algorithm is shown in Figure 1. To implement this procedure for a given model it is necessary to define bounds for input parameters and model outputs (e.g., steady states, or dynamic behavior). If bounds cannot be defined empirically, feasible ranges of parameter values can be asserted from physiological knowledge or theoretical considerations (e.g., the tissue concentration of a species may not be known, but typical weight and water content of that tissue may be known – putting an upper limit on the species concentration).

**Figure 1:**
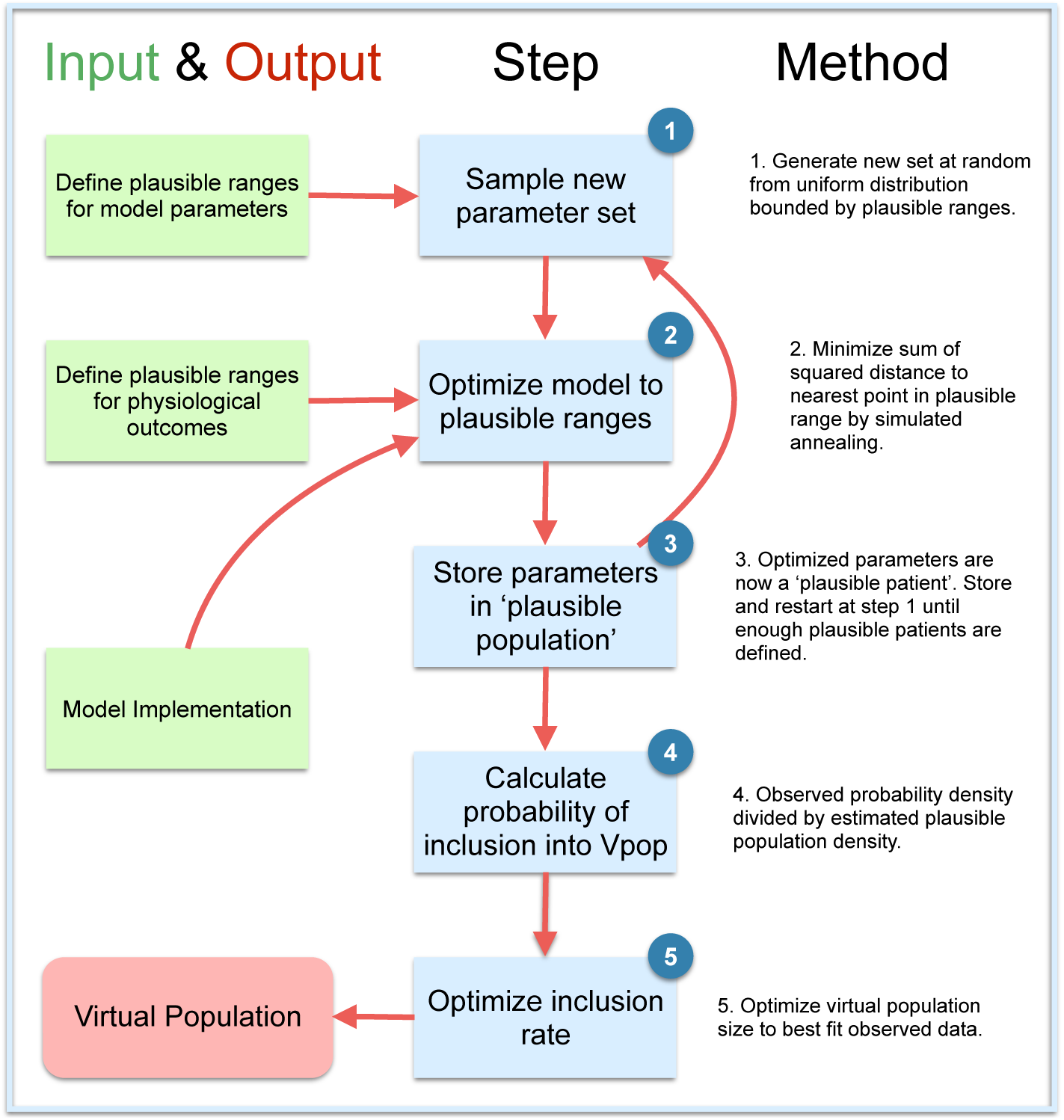
Overview of algorithm for efficient generation and prevalence-based selection of Virtual Patients. To generate virtual patients from a model, the prior information (green boxes) is used to define physiologically reasonable ranges for model outputs and parameter values. An initial parameter guess is optimized until model outputs are physiologically plausible. This is repeated multiple times to form a plausible population. A virtual population is constructed by selecting from this population with probability proportional to the prevalence in the real population relative to the prevalence in the plausible population. This selection is optimized to produce the best Virtual Population given the patients in the plausible population.

We have provided a detailed description of terminology, definitions, and the derivation of this algorithm in Table 1 and the supplementary material. Briefly, our approach is to generate a large number of ‘plausible patients.’ We define these patients as a parameter set for which every component of the model (whether it be the parameter values themselves, computed species concentrations, or combinations of these that are experimentally measureable) falls into a biologically plausible range. From this ‘plausible population’ we can then select the virtual population such that it matches the empirical distribution of interest. This is achieved by calculating a probability of inclusion of a plausible patient into the virtual population. This probability is computed from both the empirical distribution and the density of plausible patients – see the supplementary materials for more details.

**Table 1:**
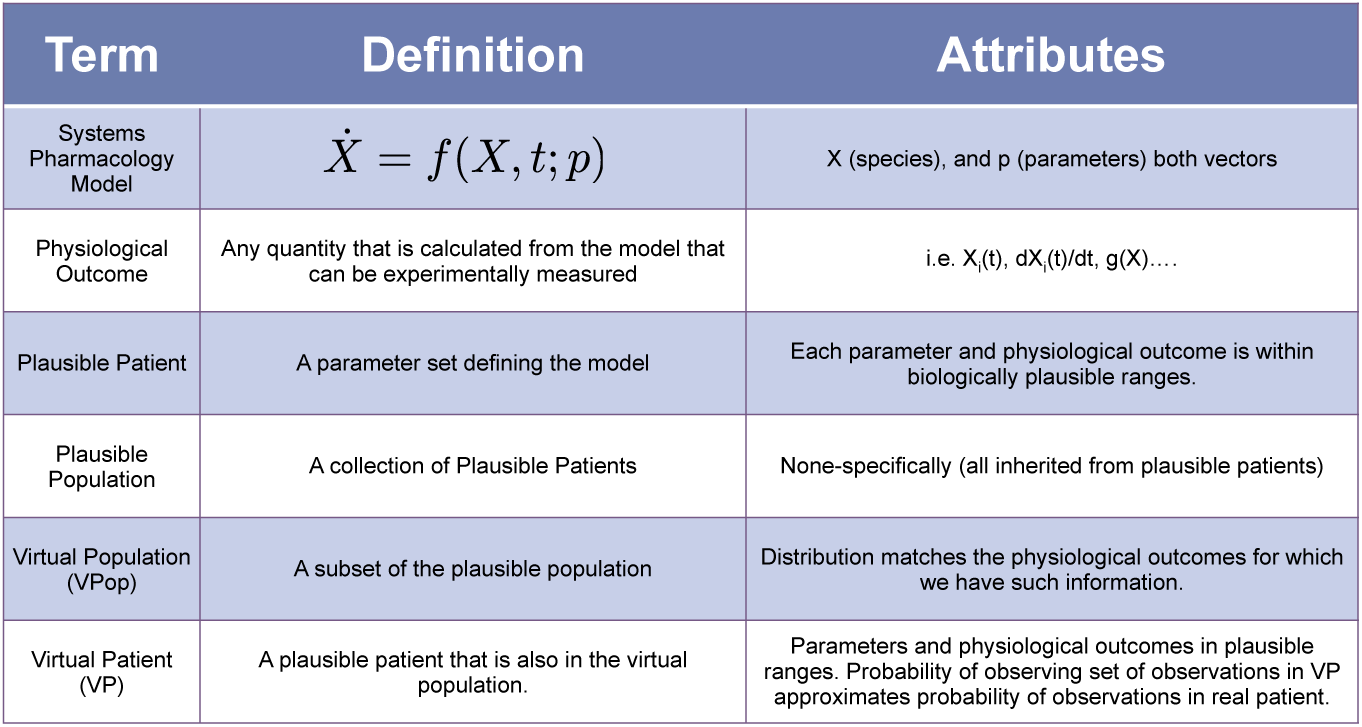
An overview of the terminology used in this paper.

An important prerequisite to this approach is the ability to generate a large number of plausible patients within the region of the empirical data. To accelerate this process we take an initial parameter guess (within the predefined bounds) and optimize this choice until the required outputs are within physiologically plausible ranges. Rather than optimize to specific points, it is more efficient to be agnostic as to where in the plausible ranges the optimization routine ends. To implement this we shift the typical cost function *f*(*p*) we would use optimizing a model to a new function, *g*(*p*), where we consider both as purely dependent on the parameter set *p*. If we constrain parameters using a number of model outputs *M_i_*(*p*), with data *d_i_* then *f* (in the simplest, unweighted case) would be

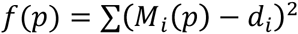

To generate plausible patients, we modify this sum-of-squared errors expression to

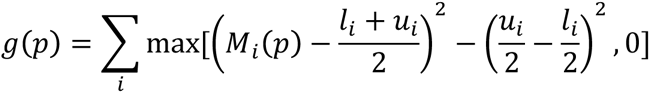

where *u_i_* and *l_i_* are the predefined plausible upper and lower bounds, respectively, for *M_i_*(*p*). This expression ensures that if *M_i_*(*p*) is in the plausible range then the contribution of the corresponding term in the expression is zero. The effect of replacing *f*(*p*) with *g*(*p*) is visualized in 2-D in Figure 2.

**Figure 2:**
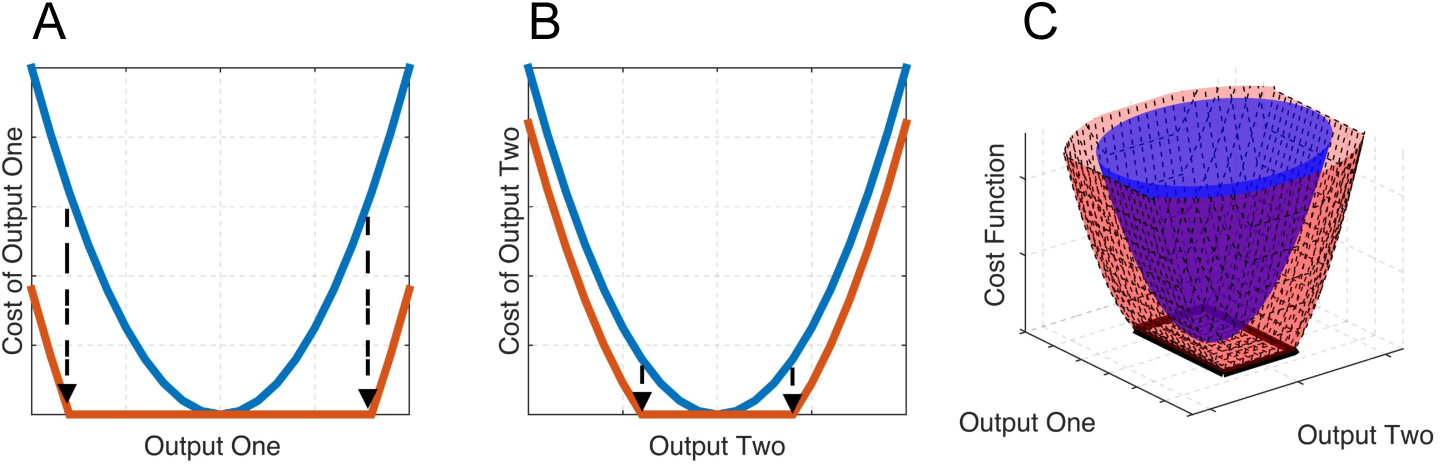
Cost-Function Transformation for convergence to plausible virtual patients. Outputs of the model contribute to the cost function to be minimized by considering the sum of squared errors (SSE) from an associated experimental observation. For each observation we define a physiologically plausible range (arrows in A and B) and shift the SSE associated with that observation so that it is zero if the model output is in this range (A and B). Combining these transformations in each dimension leads to a broader cost function that is minimized by many points, rather than one (black rectangle in C).

To test this approach we used a previously published model of cholesterol metabolism^17^. We chose this model because we could use publically available data from the NHANES database^16^ to establish the empirical multi-variate distribution for LDL cholesterol, HDL cholesterol and total cholesterol (LDL_c_, HDL_c_ and TC, respectively). Note that the distribution of these variables is well approximated by a multivariate log-normal distribution (Supplementary Figure 1). For the remainder of the paper we will describe these variables, either as model outputs or from NHANES, in log units (prior to taking the logarithm, units are mg/dl for cholesterol measures). The published version of this model does not explicitly calculate LDL_c_ or TC; instead the outputs are HDL_c_ and non-HDL_c_. From these two quantities TC is easily calculated. For full comparison with the NHANES data we introduced a new parameter to the model *k*_22_, which is simply the ratio between LDL_c_ and non-HDL_c_. Plausible ranges for the states of the model were derived from the original model publication^17^, or the NHANES data set^16^.

## Results

We generated ~300,000 plausible patients using the algorithm. As expected, the initial plausible population does not match the population-level statistics of the NHANES data (Figure 3) but covers the empirical distribution (i.e., where there are likely to be empirical observations there are plausible patients).

**Figure 3.**
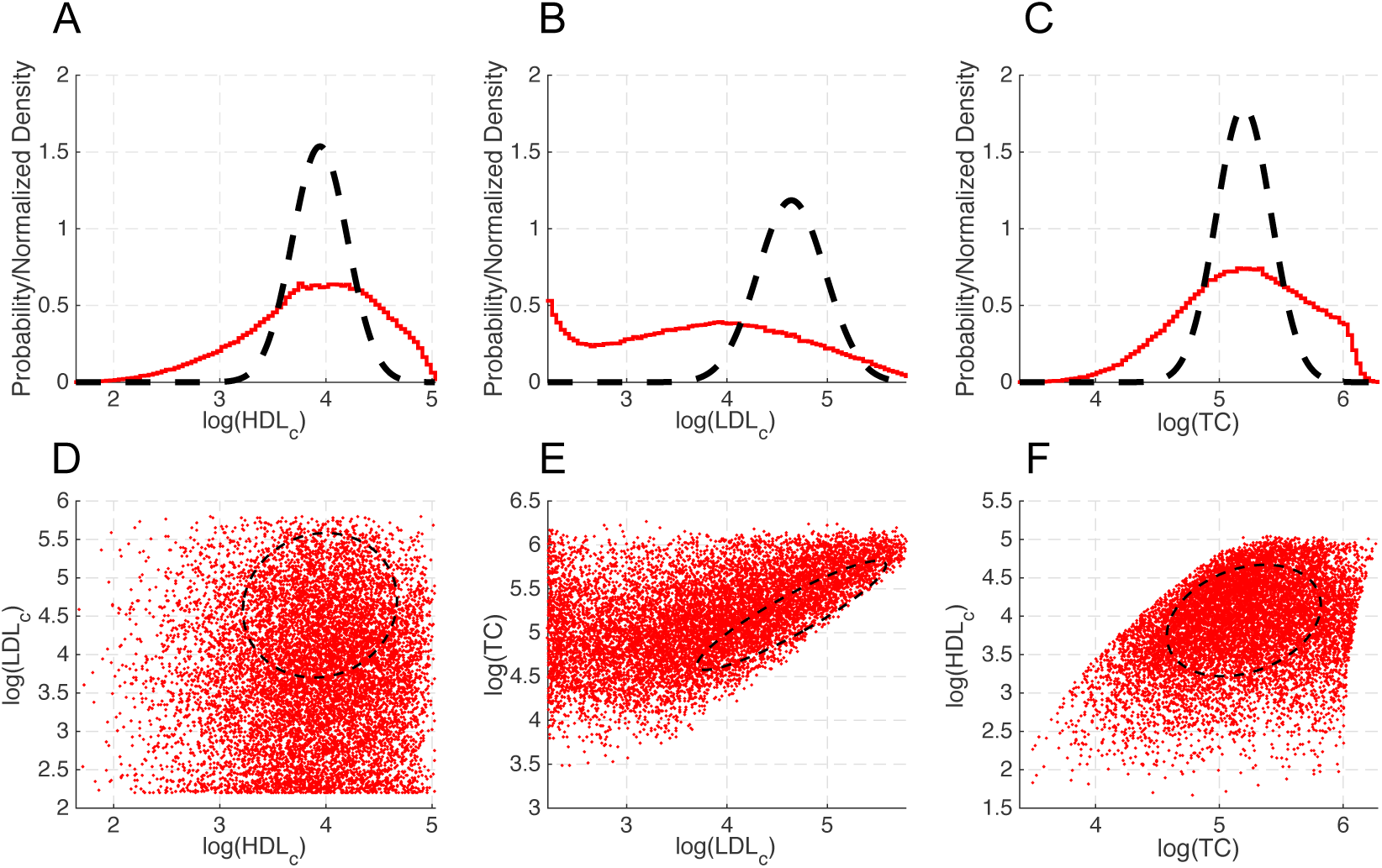
Comparison of the initial plausible population (N = 300,000) with NHANES multivariate distribution (A-C black dotted lines estimated PDF, supplementary fig 1A-C. D-F 2-D projection of the 95% confidence surface of the estimated probability density function.)

We proceeded by calculating the probability of inclusion for each ‘plausible patient’. Once calculated, we established that most of the ‘plausible patients’ are highly unlikely to be in the final distribution (Figure 4). This is due to the relative density of the plausible population to the empirical distribution. With these probabilities, only ~2% of the plausible population was selected to be in the virtual population (inset, Figure 4.). Based on examining goodness-of-fit of the distributions it appears, in this case, there is a no further value in increasing the size of the plausible population (Supplementary Figure 2).

**Figure 4:**
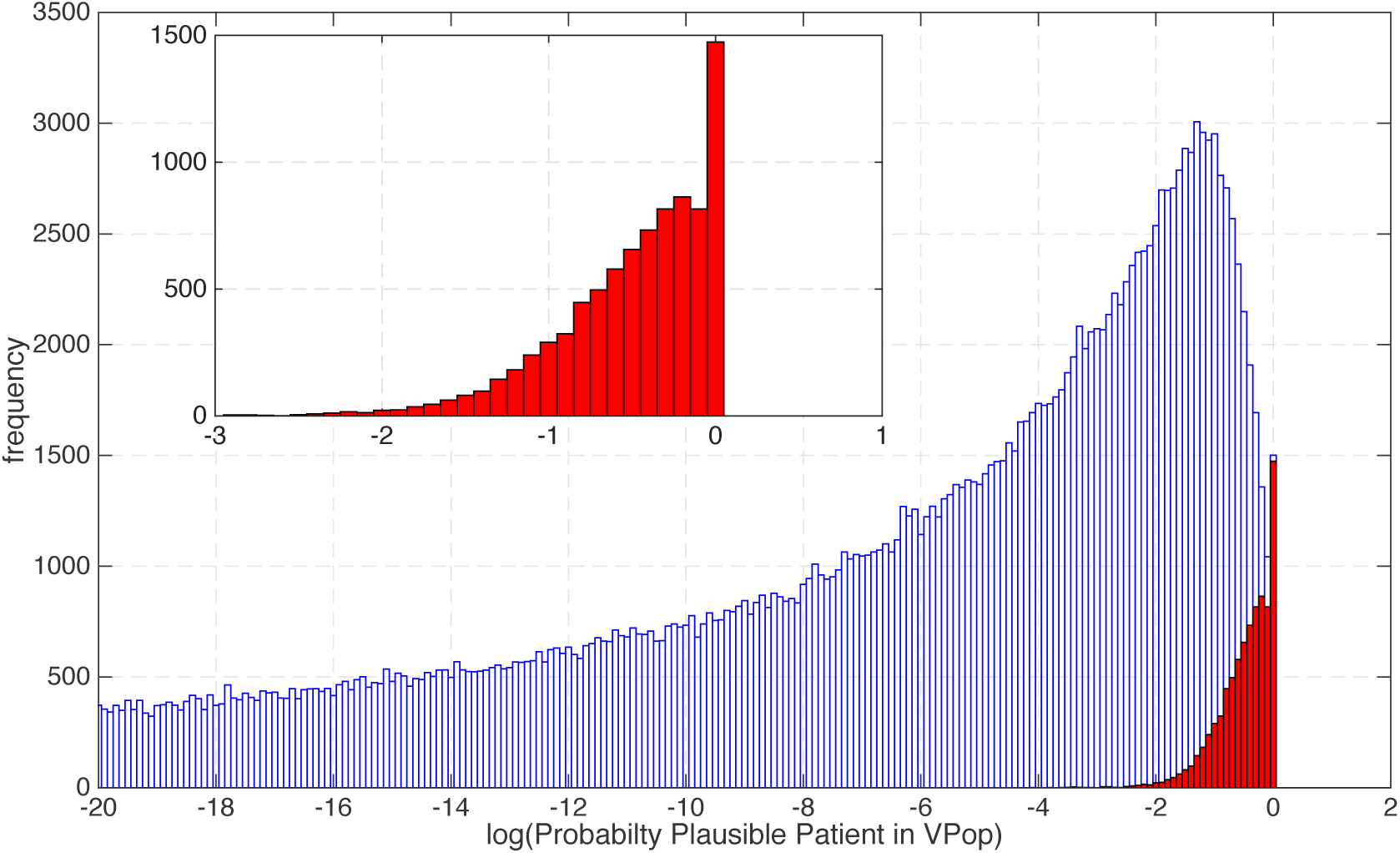
Histogram of plausible population selection probability. The probability of inclusion into the virtual population is calculated by optimized relative prevalence (see equation REF). The red histogram (main figure, and figure inset) is a virtual population that matches NHANES data, and is selected from the plausible population (blue histogram) based on displayed probability.

The distribution of an example selection fits the NHANES data well (Figure 5). The 1D histograms (when normalized for comparison with the NHANES probability density function) are indistinguishable from the data (Figures 5A-C) and the correlations between variables also match the data based on visual predictive check.

**Figure 5.**
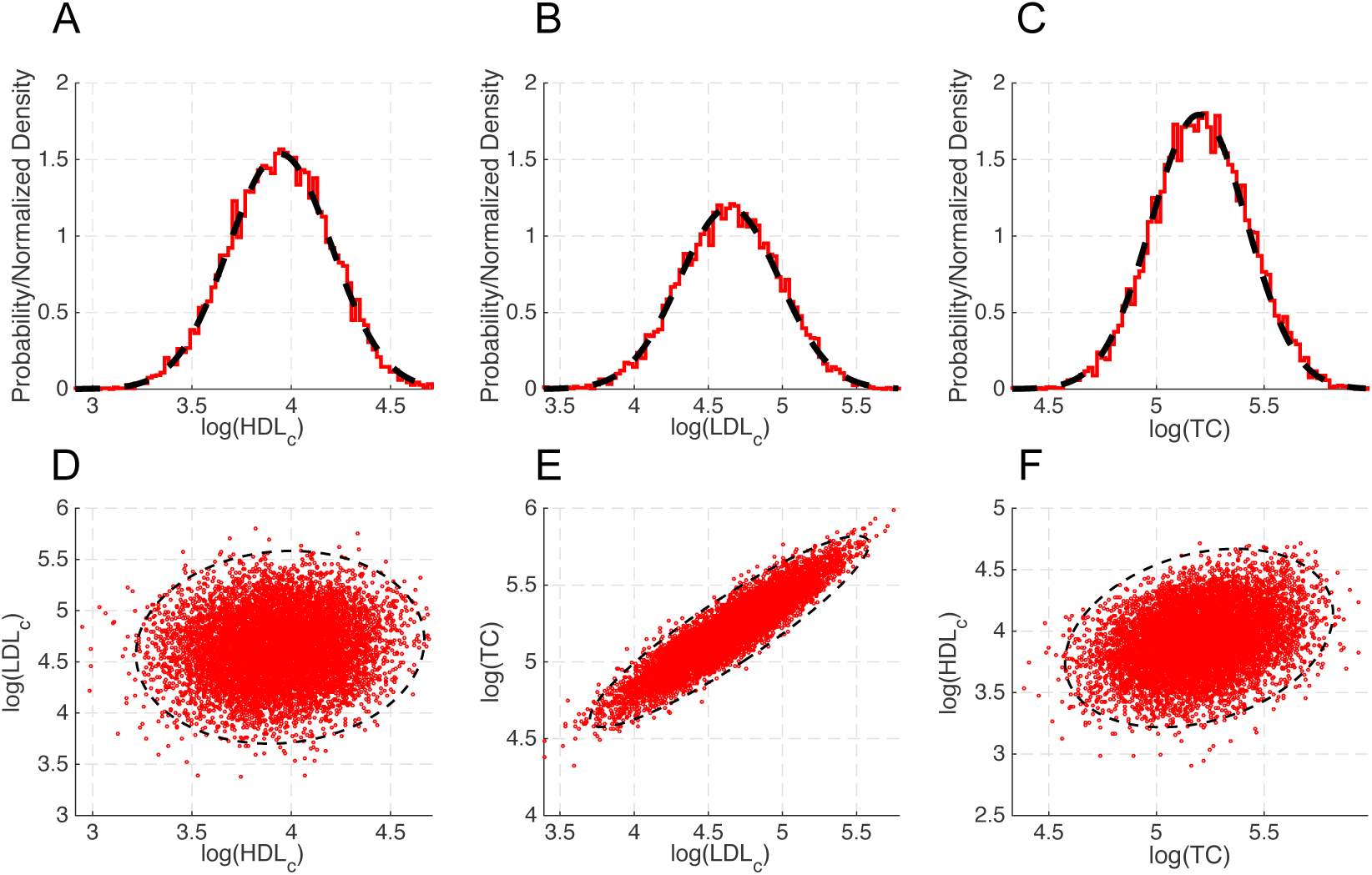
Comparison of a virtual population with NHANES multivariate distribution (dotted black lines). The virtual population (red dots and red histogram), matches the mean, variance and covariance of the multivariate experimental distribution (A-C black dotted lines estimated probability density function, supplementary fig 1A-C. D-F 2-D projection of the 95% confidence surface of the estimated PDF.)

When selecting a subset of VPs from a larger population, one concern is that the selected subset of VPs does not reflect the variability of the original ensemble, which was generated from the biologically plausible range of the parameters. Analyzing the final fitted population, we found little change in either the distribution of parameters or the correlation structure between the parameters (Figure 6 and Supplementary Figure 3). This also shows that despite the virtual population being constrained against the NHANES data the parameter values of the virtual population (Figure 6B) are only slightly better constrained than those of the plausible population. Furthermore, correlations between parameters are only slightly increased in the virtual population (Figure 6D) versus the plausible population (Figure 6C). At least in this case, constraining all outputs into realistic ranges is a more stringent constraint than selection of a virtual population.

**Figure 6 or Supplementary Figure 3:**
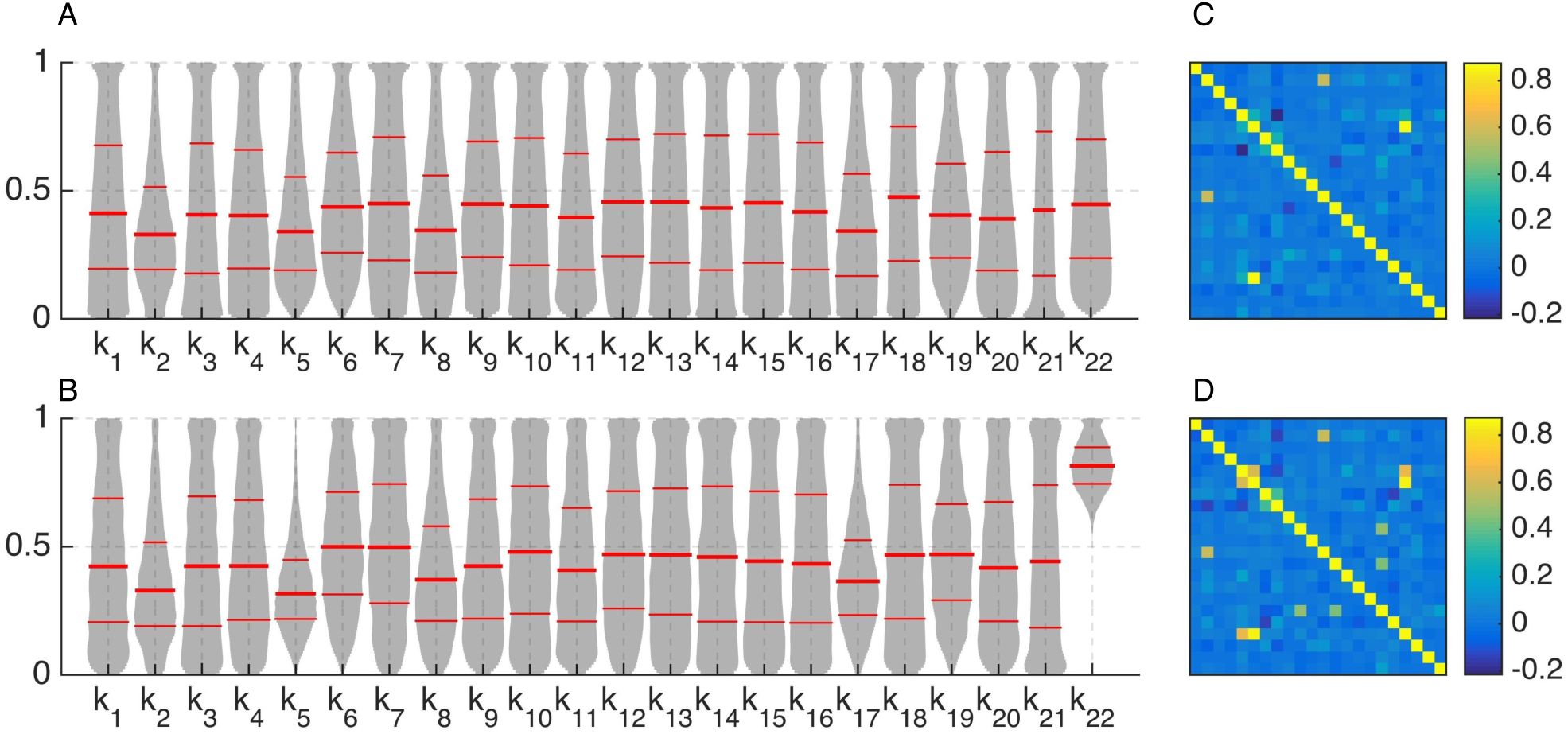
Degeneracy of the Virtual Population. Violin plots of the plausible and Virtual Populations (A and B respectively) parameter values (normalized to each parameter’s upper and lower bounds) and correlation matrix of the plausible and virtual population (C and D respectively).

## Discussion

One of the primary uses for QSP models is to prospectively simulate the effects of a dynamic perturbation (pharmacological or otherwise) in populations of interest. Due to the under-constrained nature of these models it would be difficult to have confidence in the simulation results if we simulated a single parameterization of that model, even if that set of parameters is an excellent fit to the available data. For example, imagine creating a single hypercholesterolemia VP to simulate the effect of various anti-cholesterol therapies. For the baseline characteristics of the Vpop, we have good data for the expected mean LDL and HDL (e.g., a prior clinical cohort), but we could still choose to mechanistically model hypercholesterolemia several ways using the same model. We could increase cholesterol production; decrease clearance; or apply some combination of both. Our choice of how to parameterize that VP could have significant consequences for the sensitivity of the follow-on therapy simulations. Having impaired production versus clearance of LDL could lead to differential responses to statins (production) vs. anti-proprotein convertase subtillisin/kexin type 9 (clearance). A better way is to explore the under constrained nature of these models and sample the biological uncertainty in the creation of plausible patients by varying mechanistic parameters, such as production and clearance rates, within biologically reasonable ranges. Simultaneously, we need to constrain the higher-level observables of the model based on known population distributions (e.g., the baseline characteristics of a clinical trial cohort).

An appealing aspect of the approach we outline is that since the algorithm is probabilistic once the plausible patients are generated any number of subpopulations can be selected as long as the generated patients reasonably cover the full range of the (sub)-population. Additionally, for any particular population, any number of VPops can be re-selected to bootstrap the sensitivity of the model predictions to the choice of VPop.

Achieving an acceptable fit to the data distribution is only possible if the plausible patients densely cover the range of the observables. This method should be computationally tractable for many models. However, if we have higher-order density functions, we will likely require additional gains in efficiency, above and beyond what is presented here, in methods for generating sufficient plausible patients. One potential avenue, for a future iteration of this algorithm, may be to use methods which follow a directed search through the parameter space, such as using Markov Chain Monte Carlo (MCMC) algorithms^18-20^. Although, it should be noted that this algorithm is a hybrid approach because the simulated annealing step to generate a plausible patient step is essentially an MCMC method. The advantage of this approach is it generates plausible patients (and hence virtual patients) independently – which is critical for the purpose of making a virtual population. Also, once the plausible population is established new virtual populations, suitable for new applications, can be selected.

As an introduction to this algorithm, we demonstrated how to generate a VPop that matches the baseline characteristics of a population or clinical cohort; however, in practice, a dynamic model should be constrained additionally against as many in-scope perturbation experiments as data are available. For this example, simulating changes in LDL_c_ and TC to standard-of-care lipid therapies, such as statins and ezetimibe, would likely be an important step before using the model to predict the response to a novel mechanism. Ideally, information would be available detailing the distribution of the data before and after the application of therapy (i.e., not just summary statistics). The therapeutic response can be treated as a baseline constraint for VP selection just as we used HDL_c_ and TC at baseline.

Quantitative systems pharmacology models are becoming established as a valuable component of the drug discovery and development process. Communicating their complexity and uncertainty to an interdisciplinary project team is a critical but challenging component of their utility. Virtual Populations are one tool that we have found to be successful in exploring mechanistic and parametric uncertainty in an intuitive framework that is easily understandable by most audiences. However, despite their widespread use, there are very few published methods for generating virtual patients and forming Virtual Populations. Here we have contributed an approach to efficiently generate virtual patients and construct a virtual population for which each patient is weighted equally. This method relies on the ability to generate large ‘plausible populations’, made up of plausible patients each of which is a candidate to become a virtual patient. Because a large plausible population is necessary, models that are slow to integrate (for example with dynamics across multiple time-scales) may not be good candidates for this approach. However, in cases where quantitative predictions are required and the model is amenable to thorough examination of parameter space, we have found this method to be an improvement over previous approaches.

## Supplementary Material

Consider a model *M* as a mapping from parameter space 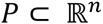to a space of observable measures, denoted 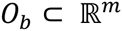:

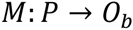

Suppose that in the real population of interest (which could be ‘all-comers’, a clinical trial population or subsets of these) that these measures, or a subset of them, have a known distribution *D*, that is *O_b_ ~D.*

Write *p* ∈ *P* as the set of *n* model parameters *p* = {*k*_1_,. *k_n_*}. Denote the parameter set *p* as a “plausible patient” if

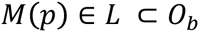

where *L* = *X*_1_× & ×*X_m_,* and *X_i_* = [*l_i_, u_i_*] is the physiologically plausible range for measurement *X_i_*. A plausible patient is one which for every measurement (when considered independently) is within physiological range, but is not a virtual patient because we have not considered whether some measurements should be correlated – hence it may be very unlikely, or impossible, to identify a real patient with similar characteristics. A *plausible population* is a collection of plausible patients, *V_pl_* = {*p_j_* | *p_j_* ∈ *L*}. As we will demonstrate, there may be a correlation structure in the plausible population due to the characteristics of the model. However, for empirical distributions there may be physiological or environmental factors external to the modeled physiology giving rise to additional relationships between physiological measurements.

To efficiently generate plausible patients we firstly bound every parameter in the model into a plausible range. For some parameters, whose values are unknown, this range will be very wide. Other parameters may be known very accurately. We use this range to generate an initial parameter guess by random sampling from a uniform distribution defined by the parameter range. Recall that, plausible patient *p_j_* is such that *M*(*p_j_*) ∈ *L*. Therefore, an initial parameter guess will likely not be a plausible patient. To find a plausible patient from an initial parameter guess we use an optimization routine seeded at the initial guess. This routine is required to converge to the space *L*, but we are agnostic as to where in *L*. In theory, we could prescribe a set of points in *L* to converge to such that the plausible population matches the data distribution *D*. In practice, it is faster to apply the following procedure and check that the generated plausible population has significant coverage of the empirical population. Consider a typical optimization problem for convergence to a set of data points *d_i_*, then the cost-function *f*(*p*) to be minimized is typically taken to be the sum of squared errors:

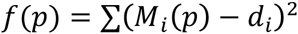

To find a plausible population we want to converge the solution to the region *L* rather than a point. To do this we will construct a new cost-function *g*(*p*). Recall *L* = *X*_1_× & ×*X_m_,* where *X_i_* = [*l_i_, u_i_*]. If we take the cost-function to be

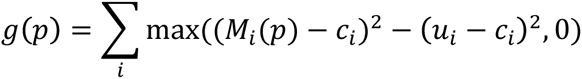

where 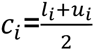, then if *M*(*p*) ∈ *L* then *g*(*p*) = 0 because this expression guarantees that if *l_i_ < M_i_*(*p*) < *u_i_* then the *i^th^* term in the sum is zero. The effect of replacing *f*(*p*) with *g*(*p*) is visualized in 2-D in figure 2.

The advantage of this approach, and in particular this cost function, is the computational cost of minimization of this function is many orders of magnitude lower than minimization of the cost-function *f*(*p*). To find multiple distinct solutions of *g*(*p*) = 0, we use repeated simulated annealing (MATLAB) with different initial guesses. The optimization can be halted once a solution to *g*(*p*) = 0 was found. Each solution we identify is then a member of the plausible population *V_pl_*.

Once we have generated *V_pl_* we can evaluate how well it matches the empirical distribution, *D*. By definition (because all measurements are in physiological range) it is plausible that *M*(*V_pl_*) could be a distributed approximately like *D*, that is 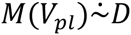, however for any relatively complex model it is highly unlikely this will be the case without refinement. We therefore need to find a subset *V_pop_* ⊆ *V_pl_* such that 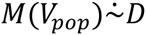, if this requirement is satisfactorily met then our subset *V_pop_* is a *virtual population* and each *p_j_* ∈ *V_pop_* is a *virtual patient.* To identify this subset we use a method closely related to the rejection-sampling algorithm, which is typically used to generate observations from a distribution where it would otherwise be difficult to do so. The distinction with the approach here is that we have a given prior distribution to select from (this is free to choose in the rejection-sampling algorithm, although does impact computation time), and we require an additional step that identifies the optimal size of the final virtual population. To illustrate this approach, consider the following.

For notation purposes, construct the selection function *S*: *V_pl_* → {0,1} as

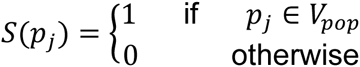

Then, from Bayes’ Theorem,

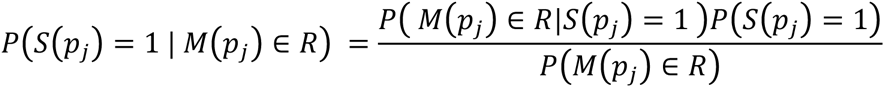

where *R* ⊆ *L* ⊂ *O_b_* is a subset of arbitrary size in the space of observable measures. Note that the left-hand side is exactly what we require: the probability a *plausible patient p_j_*, given that *M*(*p_j_*) is in the region *R*, is a *virtual patient.* Also note that, given *p_j_* is in the virtual population, we require that the probability of *M*(*p_j_*) being in the region *R* should approximately be equal to the probability of observing real data in the region *R*. That is, if *d* ∈ *D*, we can rewrite the above equation to

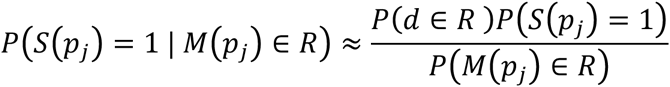

A *priori* we do not know *P*(*S*(*p_j_*) = 1), which is the probability if we pick a plausible patient at random that they are in *V_pop_.* However we do know that it is a constant for a fixed plausible population

Hence, we can write

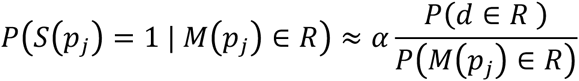

where *α* is a constant that we will fit by optimization of the *virtual population –* this controls the expected number of virtual patients in the virtual population. In the limit that the volume of the region *R*, centered around the point *r,* tends to zero.

So

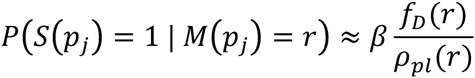

where *f_D_* is the multivariate probability density function (pdf) for the observed data *D*, and *ρ_pl_*(*r*) is an estimate of the density of the plausible population at *r.* Given that we do not normalize this estimate such that *ρ* is a pdf, the constant of proportionality is rescaled and renamed from *α* to *β*. We call this ratio ‘relative prevalence’. Intuitively, in regions where the empirical distribution is very dense selection is more likely. Conversely, where the plausible population is very dense relative to the empirical distribution selection is unlikely.

Since the plausible population may be quite sparse in some regions, a robust density estimate is required. We found that finding the spherical volume around *r*, with radius defined by the distance to the 5^th^ nearest-neighbor, *V*_5_(*r*), led to such an estimate: *ρ_pl_*(*r*) = 5/*V*_5_(*r*), which has units patients/unit volume because there are five plausible patients within the volume *V*_5_(*r*).

With this selection probability known for each plausible population, subsets matching the empirical distribution can be found by selection patients from *V_pl_* with probability as calculated above. Note that repeating this selection process will lead to a distinct virtual population.

To fit the parameter *β* we optimize the virtual population by comparison with the known empirical distribution. Goodness-of-fit is evaluated by the average (over all dimensions) univariate Kolmogorov-Smirnov test-statistic. To find the optimal value of *β* a stochastic non-gradient based method is required, because for a repeated evaluation with fixed *β* a different virtual population is selected, potentially with different goodness-of-fit to the data. To establish optimal values of *β*, we again employed simulated annealing to accomplish this. Technically, for any *r*,

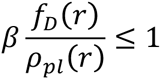

should place an upper bound on the value of *β.* However, practically we do not implement this bound and instead seek the best fit to the data (which may include plausible patients for whom this value is greater than one, which we interpret as ‘always selected’).

The success of this method hinges on generating enough plausible patients in the model that reasonably cover the empirical distribution. Otherwise, even though you can still estimate the probability of inclusion into the virtual population (provided a reasonable density estimate is possible), the expected number of virtual patients in the population (*N*. *P*(*S*(*p_j_*) = 1), where *N* is the number of plausible patients) is too low for a reasonable match to the data. Hence, failure of this algorithm is easily identified by either *N* being too small to be useful, or by a large average Kolmogorov-Smirnov test-statistic being – we have found that values greater than 0.2 tend to lead to fits that are not acceptable visually. This statistic does not assess goodness-of-fit of the cross-correlation. We recommend either visual checks or calculation of the correlation matrix for the virtual population for comparison.

**Supplementary Figure 1.**
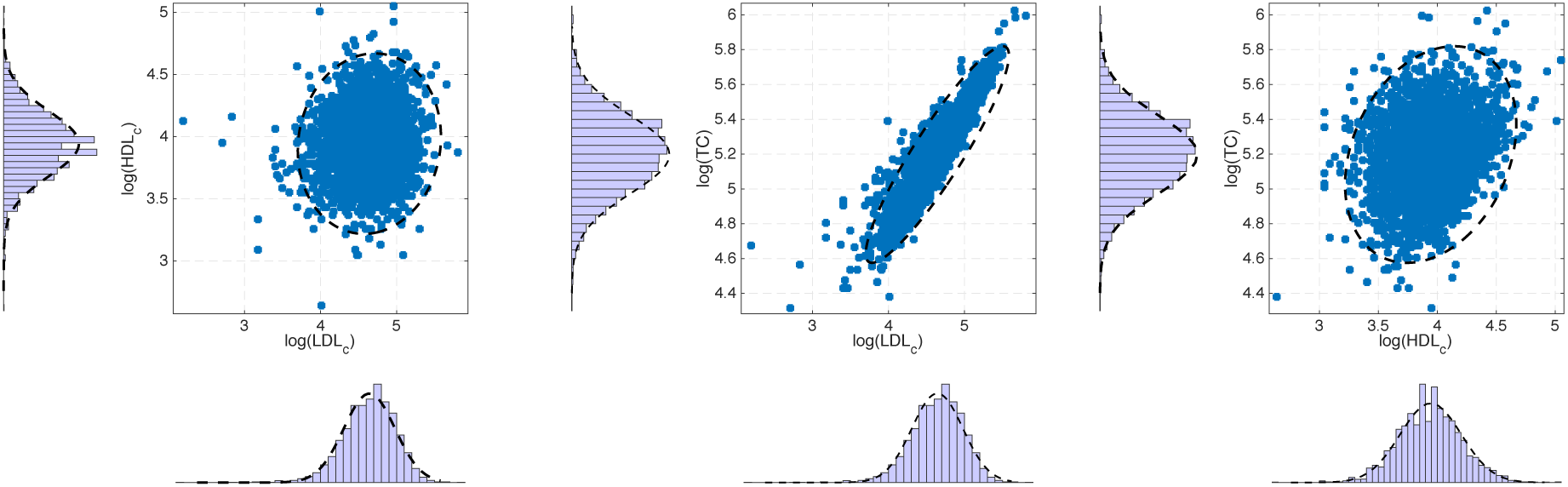
**A-C:** LDLc, HDLc and total cholesterol reported for individuals (blue circles) in the NHANES database are well described by a multivariate log-normal distribution (dotted black lines-marginal distribution fit and 2D projection of the 95% confidence surface of the estimated probability density function).

**Supplementary Figure 2.**
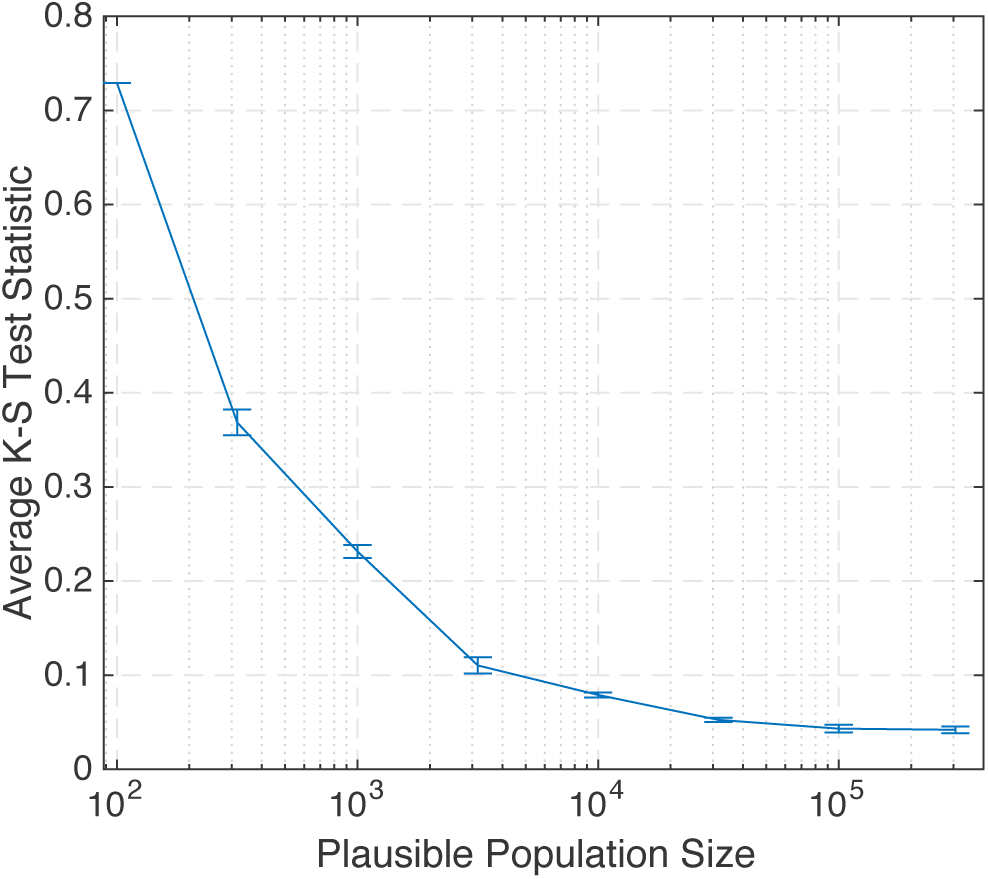
Convergence of the goodness-of-fit of the virtual population with increasing plausible population size. For this model^17^ compared with the NHANES data, a plausible population of 10,000 -100,000 is required for an optimal fit to data (as assessed by the average uni-variate Kolmogorov–Smirnov test in each dimension). Error bars are the mean +/− standard deviation of this statistic for selection of random subsets from the plausible population as described in the text.

**Supplementary Figure 3.**
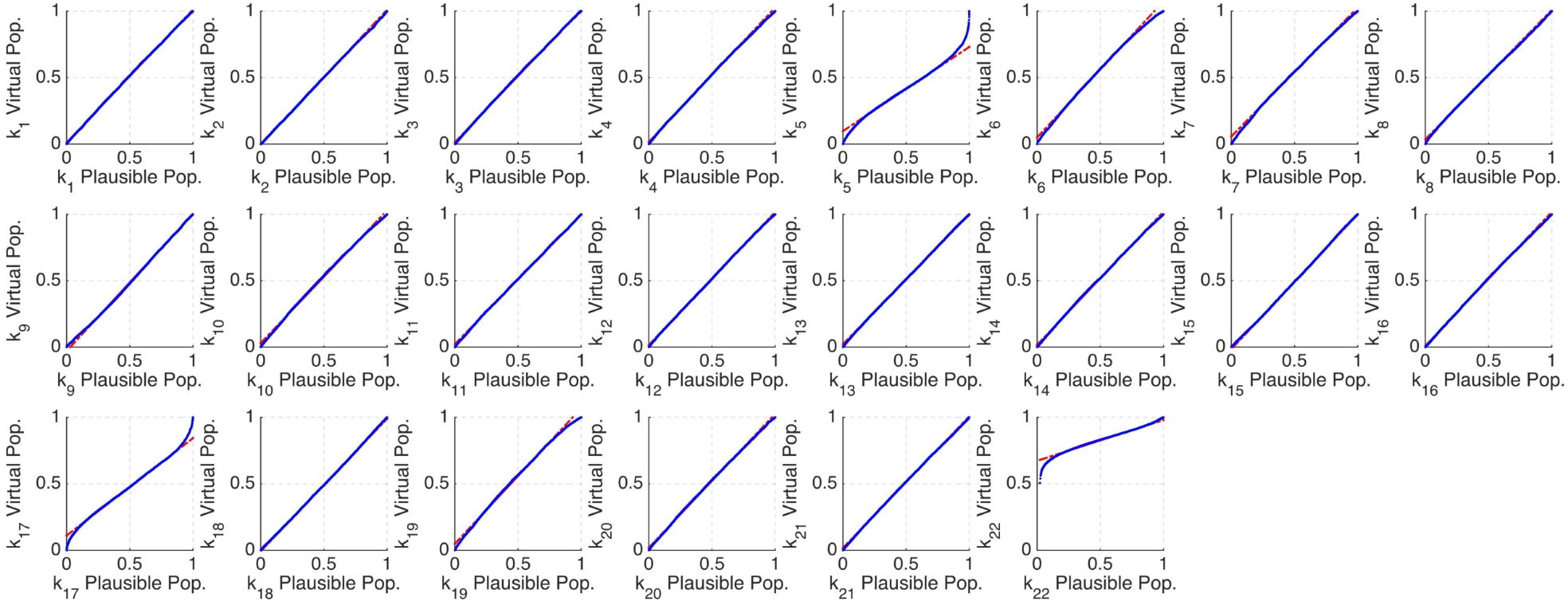
Quartile-Quartile plot of the distribution of individual parameters in the plausible population versus those in the virtual population. This shows that only several parameters are constrained by the population level fit (k_5_, k_7_, k_17_, k_22_)

